# Elucidation of the signatures of proteasome-catalysed peptide splicing

**DOI:** 10.1101/2020.04.05.025908

**Authors:** Wayne Paes, German Leonov, Thomas Partridge, Annalisa Nicastri, Nicola Ternette, Persephone Borrow

## Abstract

Proteasomes catalyse the degradation of endogenous proteins into oligopeptides, but can concurrently create spliced oligopeptides through ligation of previously non-contiguous peptide fragments. Recent studies have uncovered a formerly unappreciated role for proteasome-catalysed peptide splicing (PCPS) in the generation of non-genomically templated human leukocyte antigen class I (HLA-I)-bound *cis-*spliced peptides that can be targeted by CD8^+^ T cells in cancer and infection. However, the mechanisms defining PCPS reactions are poorly understood. Here, we experimentally define the biochemical constraints of proteasome-catalysed *cis*-splicing reactions by examination of *in vitro* proteasomal digests of a panel of viral- and self-derived polypeptide substrates using a tailored mass-spectrometry-based *de novo* sequencing workflow. We show that forward and reverse PCPS reactions display unique splicing signatures, defined by preferential fusion of distinct amino acid residues with stringent peptide length distributions, suggesting sequence- and size-dependent accessibility of splice reactants for proteasomal substrate binding pockets. Our data provide the basis for a more informed mechanistic understanding of PCPS that will facilitate future prediction of spliced peptides from protein sequences.

## Introduction

Proteasomes are multi-subunit enzyme complexes in eukaryotic cells that selectively degrade ‘unwanted’ endogenous proteins into oligopeptides. Their activity is important for protein quality control and for regulation of many intracellular processes including cell cycle progression, signalling pathways and transcription (1). The oligopeptides generated by proteasomes can be degraded to provide a source of amino acids (aas) for protein synthesis, but a subset of these peptides is translocated from the cytoplasm into the endoplasmic reticulum (ER) by the transporter associated with antigen presentation (TAP), where they may associate with newly-synthesised human leukocyte antigen class I (HLA-I) molecules (2). Peptide-loaded HLA-I molecules then traffic to the cell surface for display to CD8^+^ T cells, enabling immunosurveillance of tumours and infected cells.

The peptides comprising the cellular HLA-bound repertoire (the immunopeptidome) were classically thought to be contiguous sequences originating from self- or foreign proteins. However, following initial anecdotal reports of recognition of non-templated peptides generated by proteasome-catalysed “cut-and-pasting” of non-contiguous fragments of a polypeptide by tumour-specific CD8^+^ T cells (3-5), increasing evidence has accumulated to support the concept that proteasome-catalysed peptide splicing (PCPS) reactions comprise an additional source of peptides that can be presented on HLA-I molecules for CD8^+^ T cell recognition (6-14). In addition to generating peptides composed of fragments from within the same protein/polypeptide via *cis*-splicing, proteasomes have also been suggested to catalyse *trans*-splicing of fragments from polypeptides derived from separate proteins *in vitro* (14). However, the contribution of *trans*-spliced peptides to the immunopeptidome remains controversial and no examples of CD8^+^ T cell recognition of *trans*-spliced epitopes have been reported to date, leaving the *in vivo* significance of proteasome-catalysed *trans*-splicing unclear. Interestingly, CD4^+^ T cell recognition of HLA-II-bound epitopes composed of proinsulin peptides fused to other peptides present in beta-cell secretory granules has been implicated in the pathogenesis of diabetes (15), although the enzyme(s) responsible for the generation of these spliced epitopes were not identified.

The constitutive proteasome (CP) and the immunoproteasome (IP) both mediate PCPS reactions (6, 7), and the thymoproteasome has recently also been shown to do so (16). PCPS is initiated at proteasomal active sites by catalytic threonine residues which perform nucleophilic attack on carbonyl groups within an unfolded polypeptide chain, and can occur in either a forward or reverse sense (5, 6, 8, 17). This forms an acyl-enzyme intermediate (contained in the N-terminal portion of the final spliced peptide) which is tethered to the proteasome by an ester linkage (18). Nucleophilic attack of the acyl-enzyme intermediate by a free amine group liberated by proteasomal cleavage of a non-adjacent peptide fragment within the precursor substrate then hydrolyses the ester linkage, appending the C-terminal portion of the final spliced peptide product (4). This process is likely to be governed by inherent properties of each proteasomal active site as well as those of the constituent splice partners such as sequence, length and affinity for the substrate binding sites.

Although progress has been made in understanding proteasomal cleavage specificities during events generating non-spliced peptides (16, 19, 20), the physical constraints and biochemical preferences governing peptide splicing reactions remain undefined. Systematic analysis of PCPS reactions generating a source of peptides for HLA-I presentation has been hindered by the limited number of fully-validated spliced epitopes identified to date. The only study attempting to experimentally assign preferred ‘rules’ for PCPS was limited by its restriction to the binding preferences of HLA-A*02:01, and only considered the ligation efficiency of N- and C-terminal sequence variants of fixed lengths in a single 9-mer peptide (21).

Here, by employing a liquid chromatography tandem mass spectrometry (LC-MS/MS)-based *de novo* sequencing workflow to analyse the *in vitro* digestion products of a panel of 25 unique polypeptide substrates by the 20S CP, we have experimentally elucidated the biochemical characteristics of *cis*-PCPS reactions. We document preferred splice partner length distributions, *cis*-splicing distances and amino acid enrichments proximal to splice sites that are specifically attributable to 20S proteasomal activity and not confounded by other *in cellulo* antigen processing events, thus shedding light on the biochemical constraints inherent in both forward and reverse *cis*-PCPS reactions. Elucidation of these splicing signatures will enable refinement of a mechanistic basis for PCPS, with the eventual aim of facilitating the prediction of spliced peptides from protein sequences to enable investigation of their roles in antigen presentation and putative contribution to broader aspects of cell biology.

## Materials and Methods

### Synthetic peptides

Ten peptides derived from the sequences of NL4.3 or IIIB laboratory-adapted human immunodeficiency virus type 1 (HIV-1) strains or patient viruses, thirteen “self” peptides from different human protein sequences from the SwissProt *Homo sapiens* reference proteome and two polypeptides originating from a publicly available sORF database (22) (**Table 1**) were employed as substrates for proteasomal digestion. Peptides were synthesised by Genscript (USA) using Fmoc solid phase chemistry.

**Table 1.**
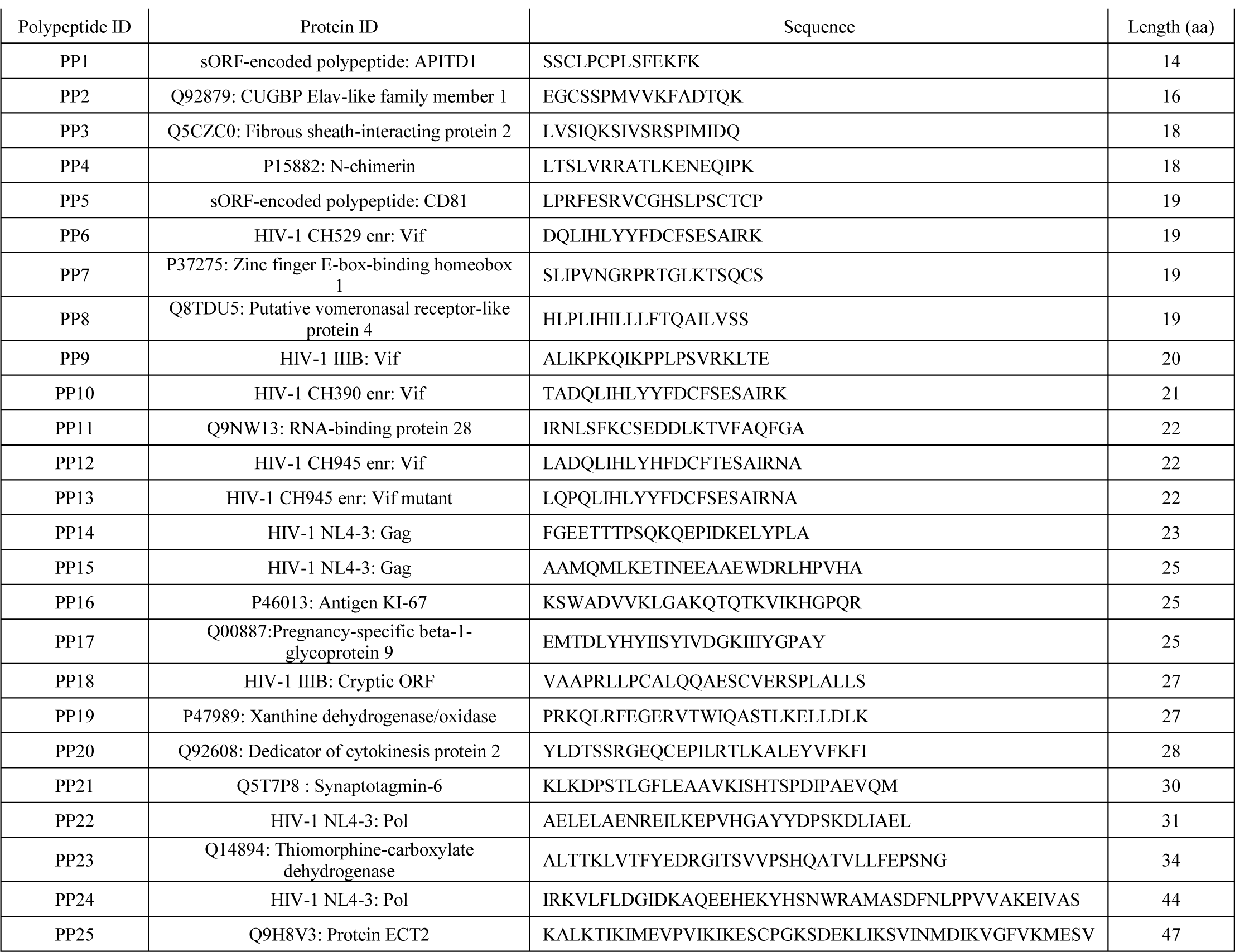
Synthetic polypeptides subjected to *in vitro* proteasomal digestion.

### *In vitro* proteasomal digests

For constitutive proteasomal digests, 5μg of synthetic polypeptide were incubated with 500ng of the 20S constitutive proteasome (CP) (Enzo Life Sciences) in 20mM 4-(2-hydroxyethyl)-1-piperazineethanesulfonic acid (HEPES) (pH 7.8), 5mM magnesium chloride, 2mM dithiothreitol (DTT) for 2h or 20h at 37°C. Control incubations (lacking the CP) were performed for each peptide in parallel. Following incubation, reactions were terminated by addition of 5μL of acetic acid, and digest material was bound to a C18 ZipTip column (Merck), eluted in 30% acetonitrile (ACN) in 5% trifluoroacetic acid (TFA), dried down and resuspended in 20μl LC-MS/MS loading buffer (1% ACN, 0.1% TFA in water).

### LC-MS/MS Analysis

1μl of each digest sample was injected onto a 3μm particle size 0.075mm x 150mm PepMap C18 trap column followed by loading onto a 2μm particle size, 75μm x 50cm analytical column on the Ultimate 3000 nUPLC system (Thermo Scientific). A gradient was applied for 30min from 8-50% buffer B (0.1% TFA in 100% ACN) in buffer A (1% ACN, 0.1% TFA in water). Peptides were introduced to the Fusion Lumos mass spectrometer (Thermo Scientific) using an Easy-Spray source at 2000V and 40°C. The ion transfer tube temperature was set to 305°C. Full MS spectra were recorded from 300-1500 m/z in the Orbitrap at 120,000 resolution with an automatic gain control (AGC) target of 400,000. Precursor selection was performed using Top Speed mode at a cycle time of 2s. Peptide ions were isolated using a width of 1.2amu, followed by trapping at a maximal injection time of 120ms, setting an AGC target of 300,000. Higher-energy collisional dissociation (HCD) fragmentation was induced at an energy setting of 28 for peptides with a charge state of 2-4, while singly charged peptides were fragmented at an energy setting of 32 at lower priority. Fragments were analysed in the Orbitrap at 30,000 resolution.

### Discovery workflow for identification of non-spliced and spliced peptides

PEAKS v8.0 (Bioinformatic Solutions) software was employed for the analysis of all LC-MS/MS datasets (.raw files). Matching against precursor polypeptide sequences was conducted without enzyme specification, using a mass tolerance precursor setting of 5ppm for peptides and 0.03Da for fragment ions. PEAKS *de novo* assisted sequencing was implemented for the assignment of non-spliced peptides derived from individual polypeptide sequences following proteasomal digest of each of the 25 precursor substrates. A false discovery rate of 5% was set using a parallel decoy database search. Following matching of spectra to respective polypeptide sequences, MS/MS spectra assigned as post-translationally modified or originating from the 20S CP were removed from the analysis, as were peptides assigned a PEAKS -10lgP score of <20, yielding a final list of non-spliced peptide sequences.

For identification of spliced peptides, a tailored bioinformatics workflow was employed similar to that previously described (9). *De novo* sequences originating from peptide spectra with average local confidence (ALC) scores of ≥50% that did not match contiguous sequences within the polypeptide precursors were considered. These were termed ‘*de novo* unmatched peptides’ (DNUPs), and only the top 5 scoring sequence interpretations for each scan were included in the analysis. As the mass:charge ratios of leucine (L) and isoleucine (I) are identical, LC-MS/MS cannot differentiate between L and I residues within peptides. Hence, PEAKS *de novo* sequencing reports all L or I residues as Ls. Therefore, for DNUPs containing a total of ‘n’ leucine residues, all permutations (2^n^) of L/I variants were computed prior to *in silico* splicing – e.g. for *de novo* sequence LTSLTLKE originating from polypeptide precursor LTSLVRRATLKENEQIPK, 2^3^ combinations of the original *de novo* sequence would be computed (LTSLTLKE, LTSITLKE, ITSLTLKE, LTSLTIKE, LTSITIKE, ITSLTIKE, ITSITLKE, ITSITIKE) and each sequence input to the splicing algorithm. If a match to a contiguous polypeptide sequence within the SwissProt proteome resulted following L/I permutations, the whole LC-MS/MS spectrum/scan was removed from the analysis.

*In silico* splicing of the resulting DNUP sequence interpretations in each scan was then implemented. Briefly, each DNUP of length ‘n’ amino acids was split into two fragments from (n-1)aa to 1aa. The (n-1)^th^ fragment was first scanned for a contiguous match across the polypeptide, and when found, its corresponding splice partner fragment was scanned for a contiguous match within the remainder of the polypeptide sequence. *Trans*-spliced peptides were omitted from the analysis. Contiguous and spliced peptides identified in control experiments involving incubation of polypeptides without the 20S CP were removed prior to further data analysis, to exclude degradation products that may not arise directly from proteasome-induced hydrolysis.

### Data analysis

Data analysis, plotting and determination of statistical significance was implemented using Graphpad Prism version 8.0. Non-spliced peptide identifications were retrieved from PEAKS PTM searches and corresponding LC-MS/MS intensity values were extracted to plot abundance comparisons. For extraction of abundance values for *de novo* sequenced *cis*-spliced peptides, matching of scan number and m/z value for a given identified *cis*-spliced peptide (obtained from the all *de novo* candidates PEAKS file) was implemented to extract corresponding intensity values from the *de novo* output. Peptide length and fragment length distributions were plotted using Prism, and peptide splicing heat maps were generated using gnuplot version 5.2.4.

### Statistical analysis

Unless otherwise stated, two-tailed unpaired Mann-Whitney t-tests were carried out to assess the statistical significance of differences between groups using GraphPad Prism v8.0. Differences were considered statistically significant at a p-value of <0.05.

## Results

### The relative proportions of unique non-spliced and *cis*-spliced peptides generated by the constitutive proteasome are dependent on precursor peptide length

To enable characterisation of non-spliced and *cis*-spliced peptides generated by the 20S CP, 25 unique polypeptides (14-47aa long, median = 22aa) were individually incubated with the CP for 2h or 20h at 37°C. Of the 25 polypeptides studied, 10 were derived from the clinically-important viral pathogen HIV-1, 13 were “self” peptides derived from human protein sequences in the SwissProt reference proteome, and 2 originated from a publicly available short open reading frame database (22) (**Table 1**). The total amino acid content of these polypeptides closely reflected that of the combined HIV-1 and human proteomes (**Table 2**). Following digestion, purification and *de novo* sequencing of proteolytic products from precursor substrates by MS, identification of both contiguous and spliced peptide products was achieved using a bioinformatics workflow (9), adapted as described in the Materials and Methods. Control digestion reactions (lacking the CP) were conducted in parallel to enable removal of background products arising from non-proteasomal polypeptide degradation or peptide synthesis artefacts.

**Table 2.** Amino acid frequencies within synthetic polypeptides and combined HIV-1 and UniProt human proteomes.

| **Amino acid <sup>a</sup>** | **Frequency in polypeptides (%)** | **Frequency in combined HIV-1/human proteomes (%)** |
| --- | --- | --- |
| A | 7.32 | 8.25 |
| C | 2.60 | 1.37 |
| D | 4.55 | 5.45 |
| E | 7.48 | 6.75 |
| F | 4.23 | 3.86 |
| G | 3.09 | 7.07 |
| H | 2.76 | 2.27 |
| I | 7.64 | 5.96 |
| K | 8.13 | 5.84 |
| L | 11.22 | 9.66 |
| M | 1.63 | 2.42 |
| N | 1.79 | 4.06 |
| P | 6.18 | 4.7 |
| Q | 4.39 | 3.93 |
| R | 4.23 | 5.53 |
| S | 8.29 | 6.56 |
| T | 4.88 | 5.34 |
| V | 6.02 | 6.87 |
| W | 0.65 | 1.08 |
| Y | 2.93 | 2.92 |

<sup>a</sup> Amino acids are denoted using single letter code

To assess the degradation of each polypeptide, we compared the in abundance (as determined from LC-MS/MS intensity values) of the precursor substrates digested by the CP at the 2h and 20h timepoints, and expressed these as a ratio of the abundance of undigested precursor (incubated without CP) at 2h. It was observed that overall, the undigested precursors underwent a significant decrease in abundance from 2 to 20h (**Supplementary Figure 1A**), indicating that saturation of available cleavage sites was not achieved at the 2h timepoint in our study. We also assessed the abundance of the shortest non-spliced peptides (5-8-mers) derived from each polypeptide substrate between the 2h and 20h time points. Here, we observed a significant increase in the median abundance of the shortest products at 20h, showing that more efficient substrate degradation was achieved by this timepoint (**Supplementary Figure 1B**). In addition, we observed a 4.5% increase in diversity of non-spliced peptides at 20h (**Supplementary Figure 1C**). However, the 2h timepoint was selected for analysis in order to minimise potential proteasomal re-entry of cleavage products and ensuing *trans*-splicing events, which could confound characterisation of *cis*-splicing products.

**Figure 1.**
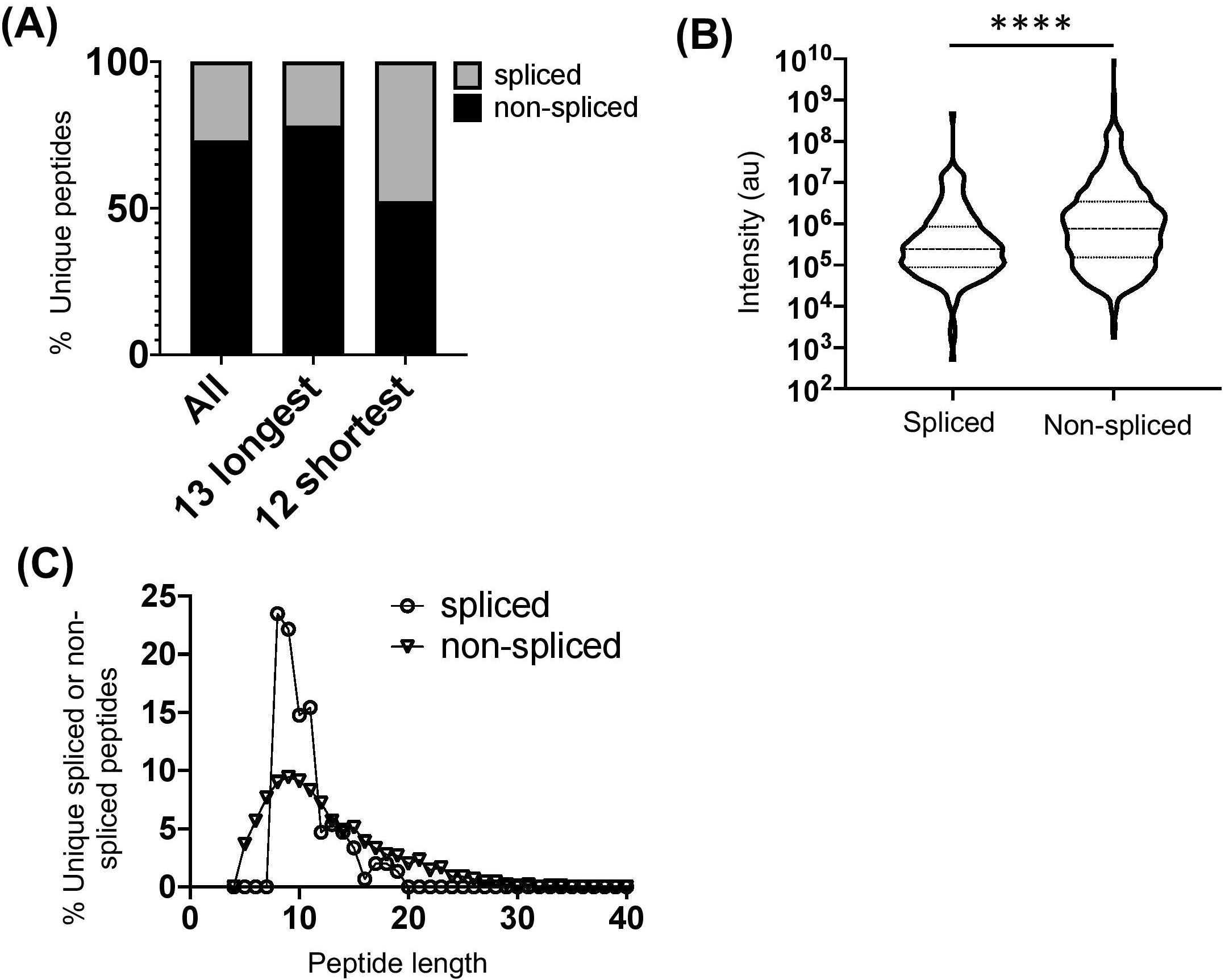
Diversity, abundance and peptide length distribution of proteasome-derived spliced and non-spliced peptides. (A) Proportion of unique spliced and non-spliced peptides following a 2h *in vitro* digestion of 25 self- and HIV-1-derived polypeptides (**Table 1**) by the constitutive proteasome. Proportions of spliced and non-spliced peptides within all unique peptides (n=1,656), unique peptides originating from only the 13 longest polypeptide substrates (n=1,337) and unique peptides originating from only the 12 shortest polypeptide precursors (n=319) are shown. (B) Violin plots showing abundance of all unique spliced and non-spliced peptides as measured by LC-MS/MS intensity values. Median and quartile abundance values are indicated. A non-parametric unpaired Mann-Whitney t-test was used to determine whether abundance values differed between groups. **** P<0.0001. (C) Length distributions of unique spliced (n=135) and non-spliced (n=900) peptides generated from within polypeptide substrates following a 2h proteasomal digest.

The full repertoire of unique spliced and non-spliced peptides generated following a 2h proteasomal digest is presented in **Supplementary Tables 1-2**. Overall, we observed a total of 1,212 unique non-spliced (73.2%) and 444 *cis*-spliced (26.8%) peptides (**Figure 1A**). The relative proportions of spliced and non-spliced peptides within all unique peptide products at the 2hr timepoint were mirrored at the 20h timepoint, where 1,270 unique non-spliced (73.1%) and 469 *cis*-spliced (26.9%) peptides were observed (**Supplementary Figure 1D**), indicating that spliced peptide generation was not under-estimated by consideration of the earlier 2h timepoint. Interestingly, however, although non-spliced peptides predominated overall, with spliced peptides constituting only approximately a quarter of the unique peptides generated, spliced peptides constituted almost half (47.6% at the 2hr timepoint) of all unique peptides generated following digestion of the 12 shortest polypeptide precursors (14-22aa) (**Figure 1A**). Furthermore, within the digestion products from precursors of all lengths, when peptide fragments containing the N- or C-terminal aa of the original polypeptide and those generated from within the polypeptide substrate (that did not then contain the N- or C-terminal aa residues) were considered separately, spliced peptides constituted a markedly lower proportion of all unique peptides in the latter category, but constituted approximately half of the peptide products containing one of the original peptide termini (**Supplementary Figure 1E**). Together, these results suggest that a substantially greater diversity of non-spliced than spliced peptides may be generated as long polypeptides derived from ubiquitinated protein substrates are digested by the proteasome within cells.

Comparison of the relative abundance (based on LC-MS/MS intensity values) of the spliced and non-spliced peptides generated from all 25 precursor polypeptides following an *in vitro* 2h proteasomal digest revealed that the median abundance of the non-spliced peptide products was significantly greater than that of the spliced peptide products (**Figure 1B**), suggesting that, at least *in vitro*, the relative efficiency of generation of non-spliced peptides is greater. Notably, the relative abundance of non-spliced peptides containing one of the terminal aas of the precursor polypeptide was significantly higher than that of peptide products generated from within the precursor (**Supplementary Figure 1F**). This is likely because non-spliced peptides containing the N- or C-termini of parental polypeptides can be generated by just a single excision event, and if higher concentrations of these peptide fragments are present they may then be preferentially incorporated into splicing events *in vitro*. However, no significant difference was observed in the median abundance of terminal amino acid-containing spliced peptides and those generated solely from within the substrate. Thus, to limit potential bias introduced in determination of PCPS signatures introduced by the differential frequencies of amino acid residues comprising the start and end of each polypeptide precursor (**Table 1**), both non-spliced and spliced peptides containing the precursor N- or C-terminal amino acid residues were excluded from subsequent analyses. Most of the splicing parameters described below remained consistent irrespective of whether analyses were performed with or without these, and any minor differences observed are noted.

### *Cis*-spliced peptides follow a narrower length distribution than non-spliced peptides

Comparison of the length distributions of *cis*-spliced and non-spliced peptides revealed that PCPS reactions generated peptides with a narrower length distribution than hydrolysis events generating non-spliced peptides (**Figure 1C**). For spliced peptides, 90% of PCPS reactions generated products that were 14aa or smaller. In contrast, the 90^th^ percentile length for non-spliced peptides was 20aa (**Figure 1C**). When both spliced and non-spliced peptides containing one of the terminal amino acids of the precursor were included, the respective 90^th^ percentile length values were 17aa (for spliced) and 22aa (for non-spliced) peptides (**Supplementary Figure 2A**). To demonstrate that this distribution was not biased by the median length (22aa) of the precursor polypeptides studied, the length distributions of *cis*-spliced and non-spliced peptides derived from just the 5 longest precursors (30-47aa) were considered. Here we observed very similar trends, where 90% of spliced peptide products were 17aa or shorter, while the corresponding 90^th^ percentile for non-spliced peptides was 24aa (**Supplementary Figure 2B**). Overall, our results thus indicate that *cis*-PCPS reactions are typically biased towards the generation of a larger proportion of shorter peptides than canonical cleavage events giving rise to non-spliced peptides. Hence, we next sought to address whether constraints such as splice partner length or other physical and biochemical factors governing splicing reactions could be identified, and whether these differed between forward and reverse *cis*-PCPS reactions.

**Figure 2.**
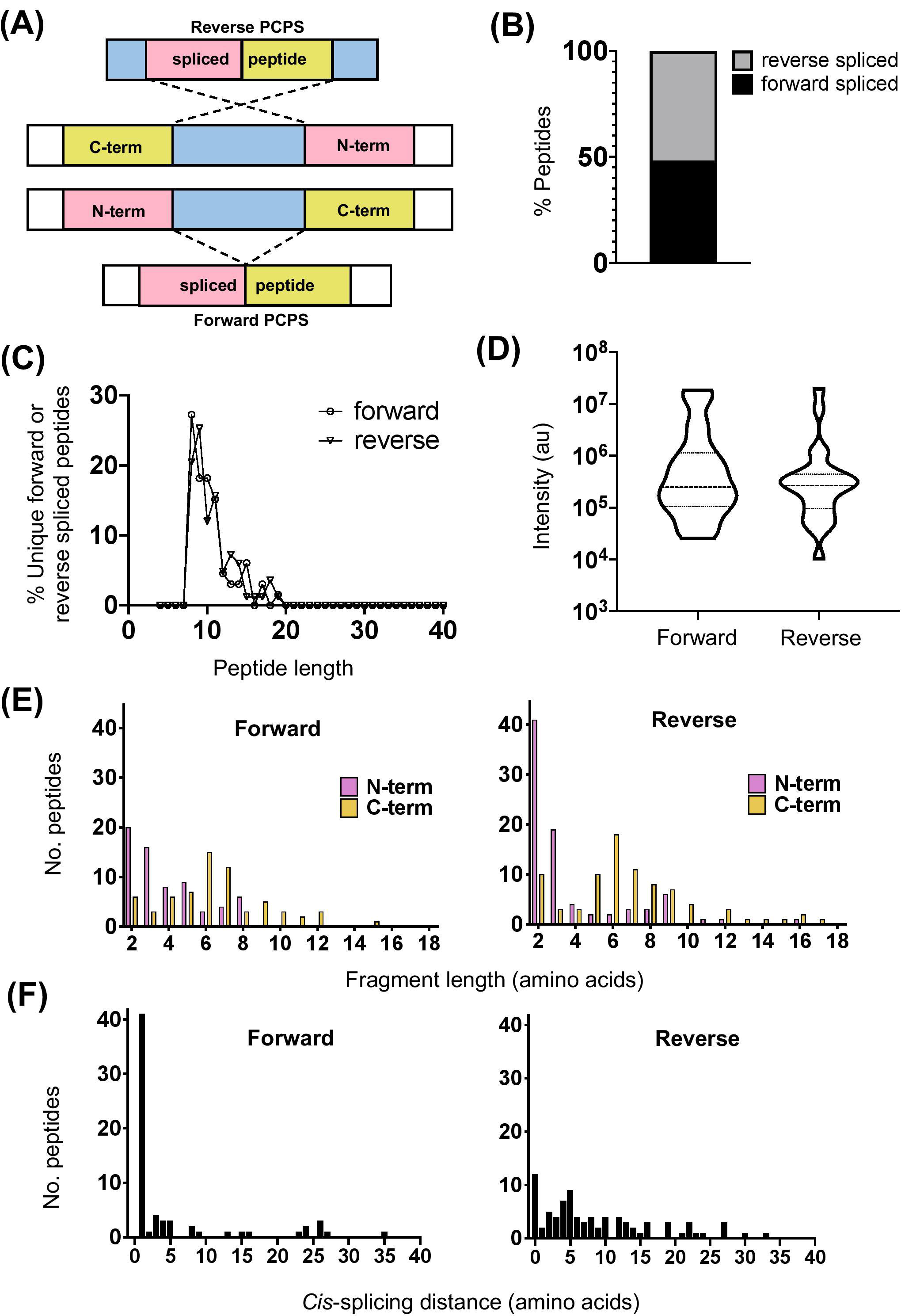
Physical properties of forward and reverse PCPS reactions. (A) Schematic depicting forward and reverse PCPS and indicating the N- and C-terminal fusion partners. (B) Relative proportions of forward and reverse spliced peptides among all unique spliced peptides (n=135) identified following 2h *in vitro* digestion of 25 self- and HIV-1-derived polypeptide sequences by the constitutive proteasome. (C) Length distributions of forward (n=65) and reverse (n=70) spliced peptides. (D) Relative abundance of forward and reverse spliced peptides. Median and quartile abundance values are indicated. A non-parametric unpaired Mann-Whitney t-test was used to determine whether abundance values differed between groups, with P<0.05 as the significance threshold. (E) Fragment length distributions of N-terminal and C-terminal splice partners involved in forward and reverse PCPS reactions. (F) *Cis*-splicing distances in forward and reverse PCPS reactions.

### Forward and reverse *cis*-splicing events demonstrate unique preferences for N- and C-terminal splice partner lengths

*Cis*-PCPS reactions occur in either a forward or reverse sense as illustrated in **Figure 2A**. To characterise the physical and biochemical properties pertaining to each type of PCPS reaction, we separated the spliced peptides identified into forward and reverse spliced subsets. Forward and reverse spliced peptides constituted similar proportions of all unique spliced peptides (48.1% forward versus 51.8% reverse spliced) (**Figure 2B**). Both sets of spliced peptides also displayed similar length distributions, indicating that both reactions predominantly generated peptides of 8-15aa in length (**Figure 2C**). Furthermore, we found no significant difference in the median abundance of forward and reverse spliced peptides (**Figure 2D**), indicating that both forward and reverse spliced peptides can be generated with similar efficiency by the CP *in vitro*.

In both types of PCPS reaction, N-terminal splice reactants were found to comprise many fragments of shorter lengths (2-5aa) (80% in forward and 66.6% in reverse spliced peptides), while longer splice reactants (6-12aa) constituted a high proportion of the C-terminal fragments of the full-length spliced peptides (**Figure 2E**). When we considered only the 5 longest precursors, we found that the fragment length distributions of forward and reverse spliced peptides were not biased by the selected polypeptide lengths, as only 12/75 (16%) of the splice partners in peptides generated from these substrates were >10aa long (**Supplementary Figure 3A**). This broadly reflected the overall distribution of splice partner fragments in **Figure 2E**, where 10.4% of the spliced partners were >10aa in length.

**Figure 3.**
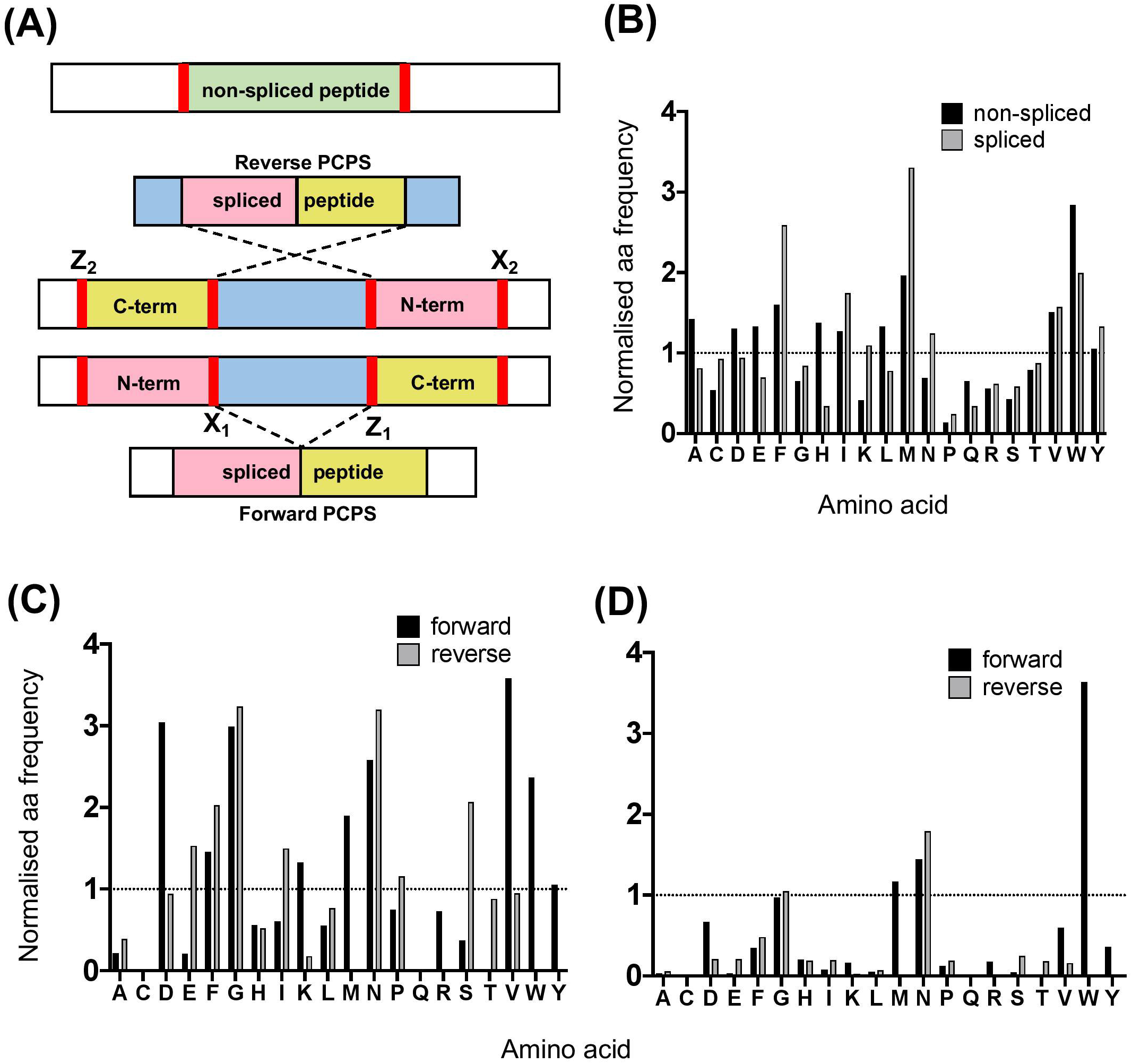
Proteasomal cleavage preferences differ between canonical hydrolysis and PCPS reactions. (A) Schematic depicting cleavage sites employed to generate non-spliced and forward and reverse spliced peptides from precursor polypeptides. X_1_ and Z_1_ denote residues at cleavage sites acting as O-acyl enzyme intermediates and nucleophilic free amine groups (respectively) in forward PCPS reactions, while X_2_ and Z_2_ denote the corresponding residues in reverse PCPS reactions. (B) Comparison of global amino acid cleavage site usage for spliced and non-spliced peptides. Amino acid frequencies are normalised relative to the overall frequency of that amino acid within all 25 precursor polypeptides. (C) Normalised frequencies of amino acid residues acting as acyl-enzyme intermediates in forward and reverse PCPS reactions. (D) Normalised frequencies of amino acid residues acting as nucleophilic free amines in forward and reverse PCPS reactions. In (B-D), amino acids are defined using single letter code.

### *Cis*-splicing events are biased towards splicing over shorter intervening distances within polypeptide substrates

Although *cis*-PCPS reactions as a whole were found to occur preferentially over shorter distances (**Figure 2F**), differences were observed in the preferred intervening *cis*-splicing distances between forward and reverse PCPS reactions. Forward spliced peptides demonstrated a more marked preference for shorter *cis*-splicing distances, with the majority of splicing events (63%) involving excision of 1-2aa and a further 18.5% occurring over *cis*-splicing distances of 3-8aa. In contrast, only 10% of reverse PCPS events occurred over a distance of 1-2aa within the polypeptide, with the proportion of PCPS events spanning a 3-8aa gap increasing to 38.6%, and 37.1% of events occurring over still longer distances. Notably, we also observed that 14.3% of reverse spliced peptides occurred with no intervening distance between splice reactants (i.e. 0 aa) (**Figure 2F**). For the 5 longest polypeptides (30-47aa) undergoing proteolysis, 10/37 (27%) of forward and 23/38 (60.5%) of reverse PCPS reactions occurred at a distance of >8aa (**Supplementary Figure 3B**). These values were greater than the proportions for each class of spliced peptide generated at a *cis*-splicing distance of >8aa observed across the entire substrate panel (18.4% for forward spliced and 37.1% for reverse spliced peptides). Thus, by using precursor polypeptides of moderate length, we may be underestimating the distances over which *cis*-PCPS can occur *in vivo* during the degradation of ubiquitinylated proteins.

### Proteasomal cleavage signatures in canonical hydrolysis and PCPS reactions demonstrate preferences for particular residues

Next, we analysed the overall frequency of cleavage after each of the 20 amino acid residues across the 25 synthetic polypeptides following both canonical hydrolysis and PCPS reactions. Global cleavage frequencies were first obtained by incorporating all cleavage sites for each set of peptides - i.e. 2 sites for non-spliced, 3 or 4 sites for reverse spliced (depending on the intervening sequence length between splice partners) and 4 sites for forward spliced peptides. These cleavage sites are denoted by red bars in **Figure 3A**. Amino acid frequencies were then normalised relative to the overall frequency of that amino acid within all 25 precursor polypeptides in order to ascertain whether enrichment of cleavage after particular residues was evident (**Figure 3B**). For non-spliced peptides, cleavage after certain amino acids (A, D, E, F, G, H, I, L, M, V, and W) was higher than would be expected given their overall frequency within the substrate precursors.

However, for spliced peptides, cleavage preferences were more restricted, with F, I, M, N, V, W and Y residues being over-represented relative to their respective frequencies in the substrate precursors. Furthermore, because PCPS is characteristically thought to involve nucleophilic attack of a stable acyl-enzyme intermediate by the free amine group of a non-adjacent amino acid sequence, we next sought to identify amino acid residues predominantly participating as either acyl-enzyme intermediates, denoted in **Figure 3A** by X_1_ or X_2_ (for forward and reverse spliced peptides respectively), or nucleophilic free amines (denoted as Z_1_ or Z_2_ for forward and reverse spliced peptides). Here, we observed differences between each reaction (**Figure 3C-D**). During reverse PCPS, E, F, G, I, N and and S predominantly acted as anchor residues for stable acyl-enzyme intermediates, while D, F, G, K, M, N, V, W and Y were typically favoured in forward splicing reactions (**Figure 3C**). Residues involved in nucleophilic attack of the N-terminal fragments were more restricted, with N specifically favoured in reverse splicing reactions and M, N and W preferred in forward PCPS (**Figure 3D**). Importantly, however, in addition to cleavage site residues, because proximal amino acids within splice partners would also dictate the affinity of each fragment for particular substrate binding pockets, we sought to determine preferred splice site signatures that may elaborate on the sequence-dependent accessibility of splice partners in forward and reverse PCPS reactions.

### Unique splice site signatures are observed in forward and reverse splicing reactions

To elucidate preferred splice site signatures, amino acid pairing combinations ± 3 residues from the fusion junction were determined. To avoid bias introduced by potential re-inclusion of proximal amino acids from multiple unique splicing events that may share the same 3 amino acids at the fusion junction (e.g. [NMD]-[KVGFVKME] and [INMD]-[KVGFVKM]), only one of the splicing events was considered to deduce the biochemical characteristics of N- and C-terminal splice partners preferentially accommodated within substrate binding pockets during PCPS (**Figure 4A**). Here, distinct patterns were observed for forward and reverse PCPS reactions (**Figures 4B** and **C**). In forward PCPS reactions, immediately proximal to the fusion junction at [P1]-[P1’] we observed preferred [P1]-[P1’] pairings (e.g. [V]–[K/N/M/W], [D]-[K], [N]-[D] and [K]-[P]) (**Figure 4B**). Conversely, in reverse PCPS reactions, we observed hotspots at [E]-[L], [F]-[S] and [D/E/S]-[K/L] at the [P1]-[P1’] position that were not evident in forward PCPS reactions (**Figure 4C**). While most acyl-enzyme intermediate and nucleophilic free amine residues participating in both forward and reverse PCPS reactions at the [P1]- [P1’] position corresponded to the dominant cleavage events (depicted in **Figure 3B**), residues from ‘minor’ cleavage sites (such as P and S) were also over-represented at [P1]-[P1’], indicating that PCPS is not entirely dictated by the efficiency of cleavage events producing the constitutive splice reactants, but is also sequence-dependent. Indeed, further differences between forward and reverse PCPS reactions were observed at [P2]-[P2’] and [P3]-[P3’] positions (**Figures 4B** and **C**), which will also determine the biochemical suitability of reactants for proteasomal substrate binding pockets.

**Figure 4.**
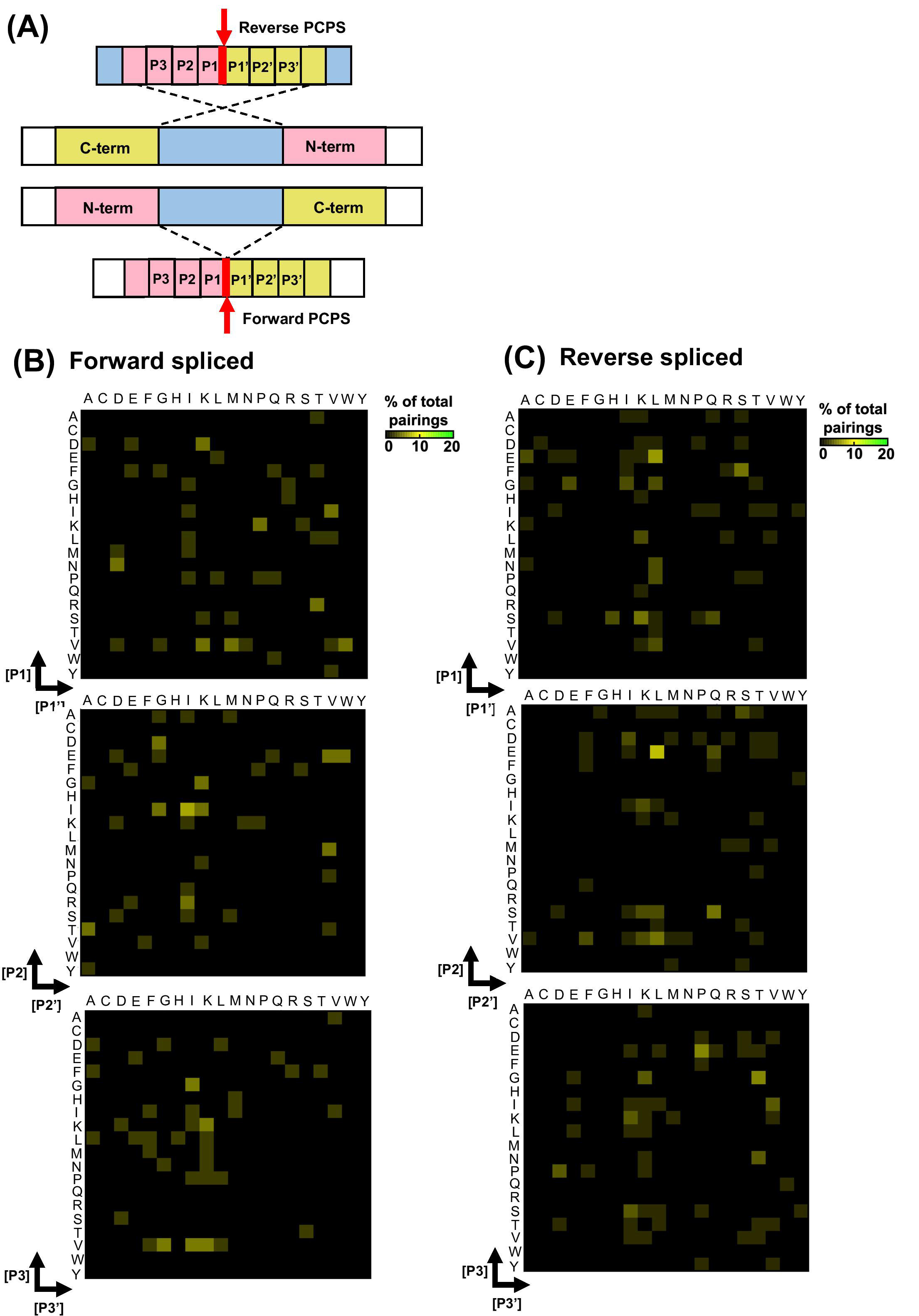
Forward and reverse PCPS reactions display unique splice site signatures. (A) Schematic depicting forward and reverse PCPS. Red arrows indicate fusion sites. Amino acid pairings proximal to the fusion site are labelled P1/P1’, P2/P2’ and P3/P3’. (B) Forward PCPS splice site signature. (C). Reverse PCPS splice site signature. In (B) and (C), heat maps show frequencies of amino acid residue pairings at ± 1, 2 or 3 aa adjacent to the fusion site. Scale represents the frequency of each amino acid pairing as a percentage of the overall pairings, and amino acids are defined using single letter code.

## Discussion

Despite over a decade of research, the properties governing PCPS are poorly understood. By employing a tailored MS-based *de novo* sequencing approach to identify peptides generated by proteasomal splicing *in vitro*, our study has thus provided novel insight into both the steric length constraints and biochemical preferences of PCPS mediated by the CP. Recent studies using MS-based approaches to investigate the characteristics of antigen processing by proteasomal isoforms have primarily focused on identification of peptides generated by canonical cleavage events (20). The development of more complex bioinformatics approaches arising concomitantly with dramatic improvements in the sensitivity of detection of MS-based technologies has yielded opportunities to more accurately and comprehensively investigate the repertoire of spliced peptides through both database-dependent as well as database-independent methodologies (9, 12, 14, 16). However, systematic analysis of PCPS has been hampered by the limited number of spliced peptides that have been experimentally-validated to date.

Although it has been proposed that PCPS reactions may be sequence-dependent (23), the only study attempting to define preferred sequence-dependent ‘rules’ for PCPS reactions to date has been limited by its restriction to the binding preferences of HLA-A*02:01, and only considered the ligation efficiency of N- and C-terminal sequence variants of fixed lengths in a single 9-mer spliced peptide [YLDW]-[KLPSV] (21). Here, the authors sought to gain insight into amino acid preferences at the splice site by either sequentially mutating the *O*-acyl enzyme intermediate of the N-terminal fragment (whilst keeping every other amino acid in the C-terminal fragment fixed), and vice versa for the nucleophilic free amine position. The two fragments were incubated with the proteasome and ligation efficiencies measured in fluorescence polarization assays. Here, it was found that D, N, Q and S predominantly functioned as *O*-acyl enzyme intermediates, whilst R was preferred as a nucleophilic free amine. This was also implemented in similar fashion for residues at [P2]-[P2’] and [P3]-[P3’]. However, importantly, the A*02:01 anchor residues were not varied in the study, and this may bias the affinities of splice partner fragments for the substrate binding pockets. Furthermore, no information could be gained on reverse-splicing events in this study, nor the length-dependent requirements and splicing distances over which PCPS events may be favored during the degradation of proteasomal substrates. Here, we experimentally define the biochemical signatures of PCPS by using a *de novo* sequencing-based approach to define spliced peptides generated following *in vitro* digestion of precursor polypeptides by the CP.

A caveat of this approach is that conditions *in vitro* do not precisely mirror those *in cellulo* e.g. are likely to be differences in the substrate:proteasome ratio and there is an absence of other cytoplasmic components, so it cannot be precluded that this introduces bias into the results obtained. However, we considered and took steps to minimise potential sources of bias that may be introduced by analysis of the spliced peptide products generated during *in vitro* PCPS reactions. In line with a previous study which showed that saturation of cleavage site usage was achieved at 480 minutes (20), we found that saturation of available cleavage sites was not achieved at the 2h timepoint in our study. Nonetheless, the 2h timepoint was selected for the investigation of PCPS reactions to minimise potential proteasomal re-entry of cleavage products as well as additional *trans*-splicing events, which could confound characterisation of *cis*-splicing products. An additional source of bias during *in vitro* processing of polypeptide substrates may arise due to systematic differences in products generated which may either have required only one cut or multiple cuts, and ensuing over-representation of N- or C-terminal amino acid residues from the precursor substrate occurring in splice site motifs. Indeed, non-spliced peptides containing the N- or C-terminal amino acid residue of the precursor polypeptide were generated in significantly greater abundance than those peptide products generated from within the precursor. However, this was not observed for spliced peptides, suggesting that the efficiency of generation of spliced peptides may not be predominantly dictated by the lower number of cleavages required to generate more highly abundant terminal amino acid-containing products, but rather by additional factors that may determine the efficiency of the PCPS reaction, such as length restrictions and biochemical properties of the splice partners themselves. These factors are likely to influence the affinity and optimal orientation of the precursor fragments for proteasomal substrate binding pockets. However, to limit bias introduced in the splice site signatures by inclusion of splice partner fragments containing terminal amino acids, the 135 of the original 444 spliced peptides that were solely generated from within the precursor substrates were utilized for analysis of PCPS characteristics.

One question we sought to address was the relative efficiency with which spliced and non-spliced peptides are generated by the CP. In the *in vitro* system studied here, spliced peptides were found to constitute a relatively low proportion of the products generated from within precursor polypeptides, suggesting that a greater diversity of non-spliced than spliced peptides may be generated as long polypeptides derived from ubiquitinated protein substrates are digested by the CP *in vivo*; and the abundance of the spliced peptides was also found to be significantly lower than that of non-spliced peptides (**Figure 1A-B**). Prior studies where the relative abundance of spliced and non-spliced peptides generated by other types of proteasomes was addressed have not yielded a consensus about this (7, 9), but these utilised only very small numbers of polypeptide substrates, suggesting a need for more comprehensive future analyses to clarify whether there are differences in the efficiency with which spliced and non-spliced peptides are generated by different proteasomal isoforms both *in vitro* and *in cellulo*.

To date, it has been unclear whether stringent length requirements of splice reactants are required for enhanced efficiency of PCPS reactions. We observed that *cis*-PCPS reactions typically generated a larger proportion of shorter peptides than cleavage events giving rise to non-spliced peptides, and that the majority of N- and C-terminal splice partner fragments were typically between 2-7 aa in length (**Figure 2E**). We postulate that this could be due to the nature of the substrate binding pockets and/or amino acid residues surrounding the catalytic subunits, which in turn govern the optimum lengths of splice partner fragments that can be accommodated in the correct orientation to facilitate peptide splicing. The similar splice partner fragment length distributions observed for the 5 longest polypeptide precursors (**Supplementary Figure 3A**) further suggests that steric constraints within proteasomal substrate binding pockets may limit the length of splice partners that can be accommodated in PCPS reactions. Although it has previously been shown that the minimal length of a C-terminal splice reactant required to perform nucleophilic attack on a stable acyl-enzyme intermediate was 3aa (8), no upper bound for the length of splice partners has previously been described. Based on our results, this value may typically lie between 9-12aa. However, although unlikely at the 2h digestion time point analysed here, we cannot entirely preclude that post-PCPS processing (via proteasomal re-entry and cleavage of previously spliced peptide products) may bias the fragment length distributions observed.

Data from our study also elucidate the distances over which PCPS can occur within a polypeptide substrate, and show that these distributions differ markedly between forward and reverse splicing reactions. We demonstrate that while forward splicing typically involves the excision of 1-2 aas, a larger proportion of reverse PCPS reactions occur over distances greater than 3 aas (**Figure 2F**). However, we also show that the use of precursor polypeptides of moderate length may lead to an underestimation of the distances over which *cis*-PCPS can occur, particularly with regard to reverse PCPS events (**Supplementary >Figure 3B**). Indeed, an experimentally validated fibroblast growth factor (FGF)-5-derived spliced epitope in renal cell carcinoma was experimentally shown to be generated by a transpeptidation reaction spanning 40aa (3). However, it was subsequently demonstrated that *cis*-PCPS reaction efficiencies displayed an inverse correlation with increasing intervening sequence length (6), indicating that PCPS reactions are likely to occur with greater efficiency over shorter distances.

Finally, we sought to determine both cleavage and sequence-dependent fusion preferences of splice reactants within the proteasomal chamber. As PCPS is characteristically thought to involve nucleophilic attack of a stable acyl-enzyme intermediate by the free amine group of a non-adjacent amino acid sequence (it has also been shown that certain spliced peptides may be produced within cells without the formation of a semi-stable *O*-acyl-enzyme intermediate via a condensation reaction (7)), we experimentally defined residues predominantly acting as stable *O*-acyl-enzyme intermediates and nucleophilic free amines. Although the global sequence context of individual precursor polypeptides may influence cleavage specificity, our data suggest that in addition to preferred length constraints for N- and C-terminal splice partners, certain residues may preferentially function as more stable reactive intermediates or as nucleophilic free amines in forward and reverse PCPS reactions, which may facilitate optimal positioning or stability within the proteasomal chamber to enable splicing. For non-spliced peptides, although we did not observe pronounced tryptic-like activity (cleavage after basic residues) our data are largely in accordance with results from a recent analysis of non-spliced proteasomal cleavage products derived from a set of 228 rationally-designed synthetic 14-mer peptides, where preferential cleavage after A, F, H and L residues was observed (20). Additionally, while it is established that the catalytically active subunits (β_1_, β_2_ and β_5_) of the CP predominantly initiate cleavage after acidic (D and E), hydrophobic (F, W and Y) and basic residues (K and R), respectively (18), cleavages following branched chain (L, I and V) or small neutral amino acids (A, S, T and V) also predominate (24), and were likewise observed in our study.

As with cleavage signatures, we also found differences in the enrichment of particular amino acid pairings in forward and reverse PCPS reactions by examination of splice site signatures (**Figure 4B**). These data may suggest that forward and reverse PCPS events may occur in distinct substrate binding sites within the proteasomal chamber. Interestingly, we observed that at the fusion site, in addition to amino acid residues derived from the dominant cleavage events, residues from ‘minor’ cleavage sites were also enriched at the [P1]-[P1’] position, demonstrating that PCPS is sequence-dependent, and not entirely governed by the frequency of splice reactant precursors generated by the dominant cleavage events. Importantly, inspection of splice site signatures defined following *in vitro* proteasomal digests in our study revealed that these differed from those of previously-reported HLA-I bound spliced peptides identified from MS-based immunopeptidome profiling datasets (12, 14). Previously proposed amino acid splicing preferences determined by inspection of immunopeptidomic datasets could be biased by selective pressures imposed on proteasome-generated peptide pools during subsequent steps of antigen processing/presentation, e.g. TAP-mediated peptide translocation into the ER, HLA-I binding preferences and ER-mediated peptide trimming, resulting in omission of PCPS-derived peptides (25). The repertoire of spliced peptides detected in HLA-I-bound peptide repertoires may also be restricted by the peptide length preferences of class I alleles. For instance, in our study, ∼35% of *cis*-spliced peptides generated following *in vitro* proteasomal polypeptide digestion were >12aa in length, and would thus be under-represented in HLA-I bound peptide datasets.

Taken together, our experimental characterization of PCPS reactions using *in vitro* digests of polypeptide substrates indicate certain well-defined characteristics of peptide splicing by the CP, and allows us to propose a model for peptide splicing by the CP. **Figure 5** illustrates the key features of forward and reverse PCPS reactions defined in our study. We note that in both forward and reverse PCPS reactions (**Figure 5A-C**) N-terminal fragments were usually shorter, while C-terminal splice reactants were of longer lengths. However, *cis*-splicing distances in reverse PCPS were typically much longer than in forward PCPS reactions. The etiology of this major difference is unclear, although one may postulate that steric constraints within the proteasomal binding chamber may require the N-terminal portion of the precursor to re-orientate and perform nucleophilic attack of the C-terminal end in an adjacent substrate binding pocket (**Figure 5B**), thus causing the polypeptide to re-orientate and bend within the proteasomal chamber. Longer intervening *cis*-splicing distances might thus allow for greater flexibility of the polypeptide chain. This may occur similarly in reverse PCPS reactions with no intervening *cis*-splicing distance (**Figure 5C**).

**Figure 5.**
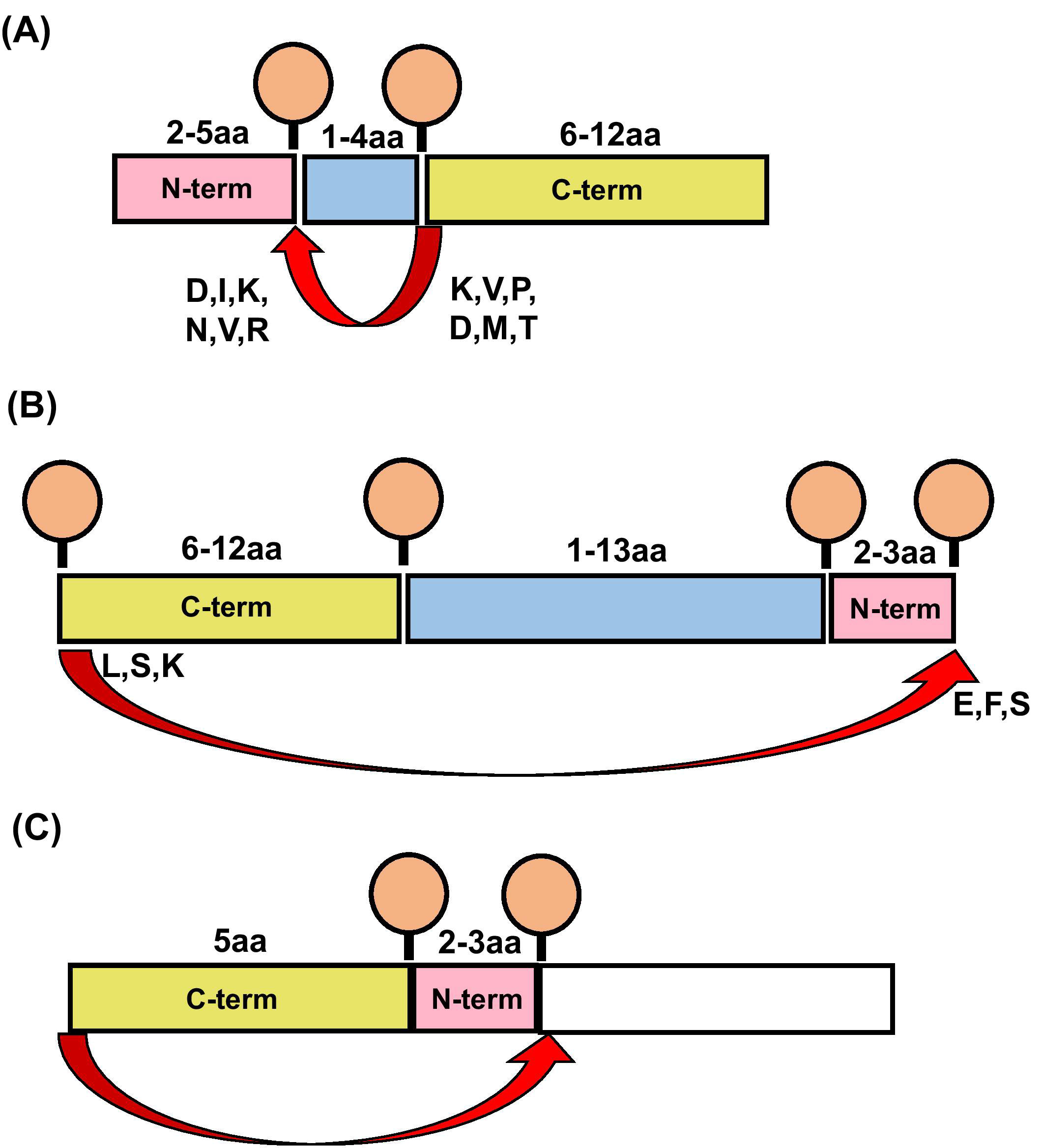
Schematics depicting characteristics of proteasome-catalysed peptide splicing. (A) Diagram illustrating the characteristics of forward PCPS events. (B) Diagram illustrating the characteristics of reverse PCPS events occurring over *cis*-splicing distances >0aa. (C) Diagram illustrating the characteristics of reverse PCPS events occurring with no intervening *cis*-splicing distance. In (A), (B) and (C), Red arrows indicate splicing events. Amino acids typically acting as *O*-acyl enzyme intermediates or nucleophilic free amines as defined in splice site signatures (**Figure 4**) are shown in single letter code at the end and start of N- and C-terminal splice partners (respectively).

Future analysis of *in vitro* peptide splicing by other proteasomal isoforms, and comparative analysis of the HLA-I bound spliced peptides generated in cells expressing different proteasomal subunits, and in immunopeptidomes generated by direct versus cross-presentation would be of interest. Importantly, the delineation of cleavage and splice site signatures reported here will facilitate a more informed investigation of the mechanisms of PCPS, and the development of algorithms to predict spliced peptide generation from protein sequences. This will in turn expedite future work to define spliced peptides derived from pathogen and tumour antigens that could be targeted for prophylaxis or therapy. In addition, a more thorough understanding of the repertoire of spliced peptides produced within cells could uncover roles of these peptides in aspects of cell biology beyond antigen presentation.

## Supporting information

Supplementary Figures 1-3 REVISED

Supplementary Tables 1 and 2 REVISED

## Author Contributions

WP, GL, TP, NT and PB conceived the study. WP designed and conducted experiments, analysed data, prepared figures and drafted the manuscript. AN and NT acquired MS data. GL, TP, NT and PB contributed to data analysis and manuscript editing. All authors read and approved the final version of the manuscript.

## Acknowledgments

The authors would like to thank Professor Benoit van den Eynde for helpful discussions about the study. This manuscript has been released as a pre-print at bioRxiv (26).

