## Supplementary Figures 1-3 REVISED for "Elucidation of the signatures of proteasome-catalysed peptide splicing"

### Supplementary Figure 1

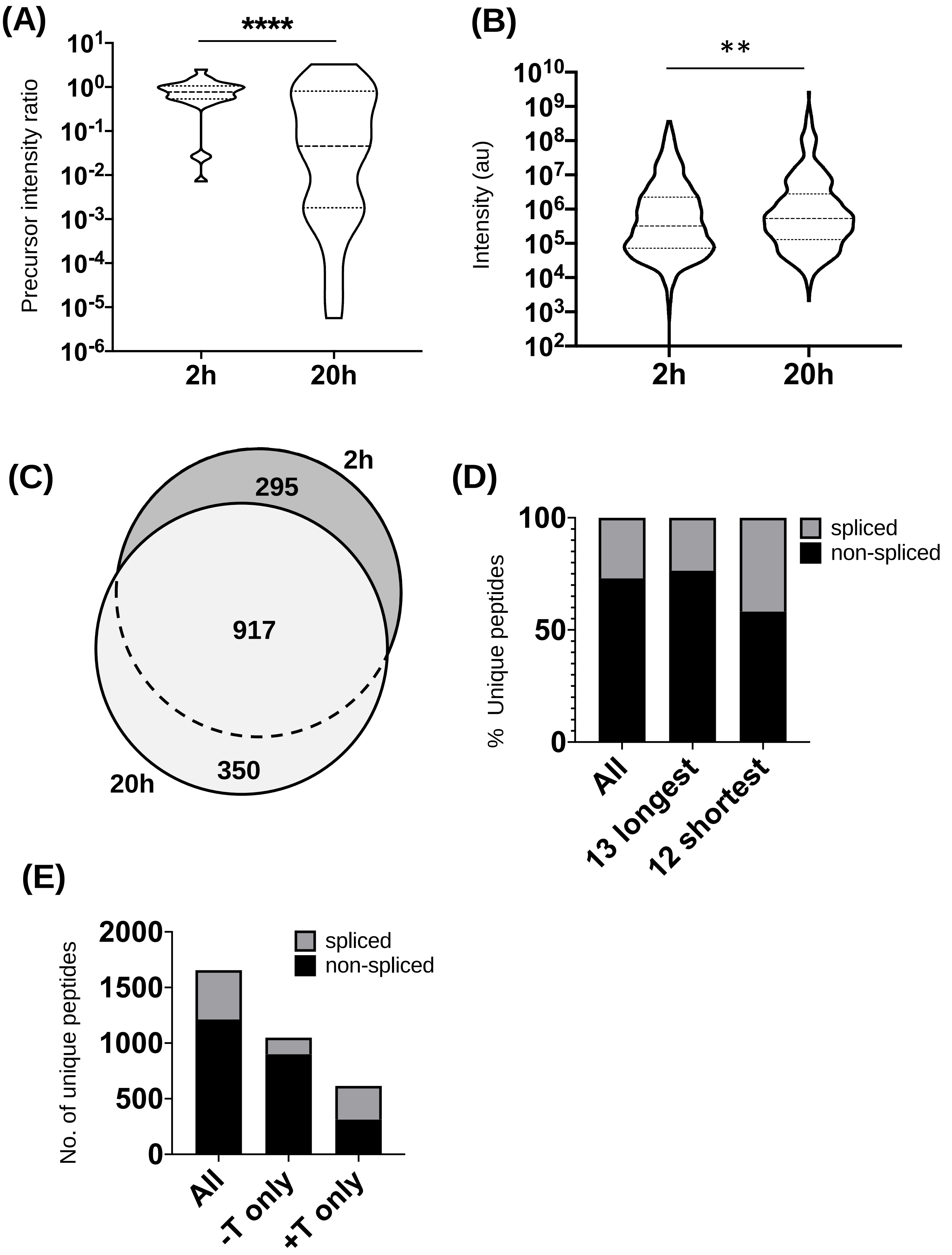

**(F)**

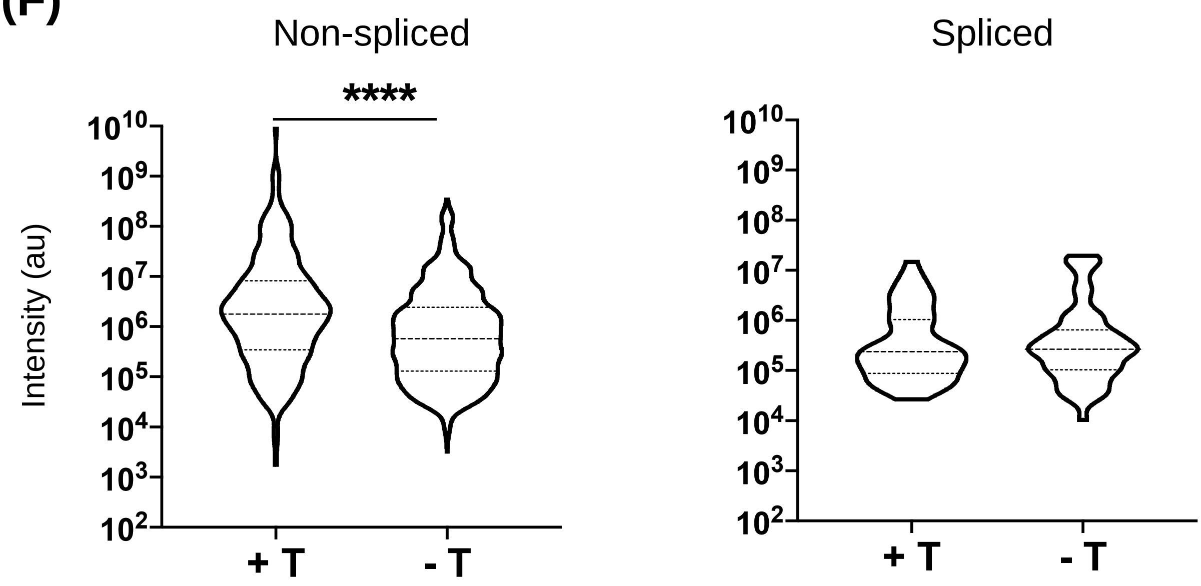


**Supplementary Figure 1. Diversity and abundance of proteasome-derived spliced and non-spliced peptides**

(A). Comparison of the abundance ratios (as quantified by LC-MS/MS intensity values) of full-length precursor polypeptides (**Table 1**) measured at the 2h and 20h timepoints relative to the undigested precursor at 2h. Median and quartile abundance ratio values are indicated. Statistical significance was calculated using a ratio paired t-test. **** P<0.001.

(B). Comparison of the relative abundance (as measured by LC-MS/MS intensity values) of total 5-8-mer peptides generated at 2h or 20h. Median and quartile abundance values are shown. ** P<0.01.

(C). Area-proportional Venn diagram illustrating the diversity of constitutive proteasome-derived non-spliced peptides following *in vitro* digestion of precursor polypeptide substrates for 2h or 20h.

(D). Proportion of unique spliced and non-spliced peptides following 20h *in vitro* digestion of 25 self- and HIV-1-derived polypeptide sequences by the constitutive proteasome. Proportions of the unique peptides (n=1,739) generated from all 25 polypeptide substrates, unique peptides originating from only the 13 longest polypeptide substrates (n=1,414) and unique peptides originating from only the 12 shortest polypeptide precursors (n=325) are shown.

(E). Number of unique spliced and non-spliced peptide products identified following 2h *in vitro* digestion of 25 precursor polypeptides by the constitutive proteasome. Numbers of all unique spliced and non-spliced peptides, those originating from within the polypeptide substrate and not containing the terminal amino acid (-T) and those containing terminal amino acid(s) of the precursor substrate (+T) are shown.

(F). Violin plots showing abundance of the spliced and non-spliced peptide products containing terminal amino acids of precursor substrates (+T) and those originating from within the polypeptide substrate (-T), as measured by LC-MS/MS intensity values. Median and quartile abundance values are indicated. ****P≤0.0001. In (B) and (F), P<0.05 was used as the threshold for significance following a non-parametric unpaired Mann-Whitney t-test. .

### Supplementary Figure 2

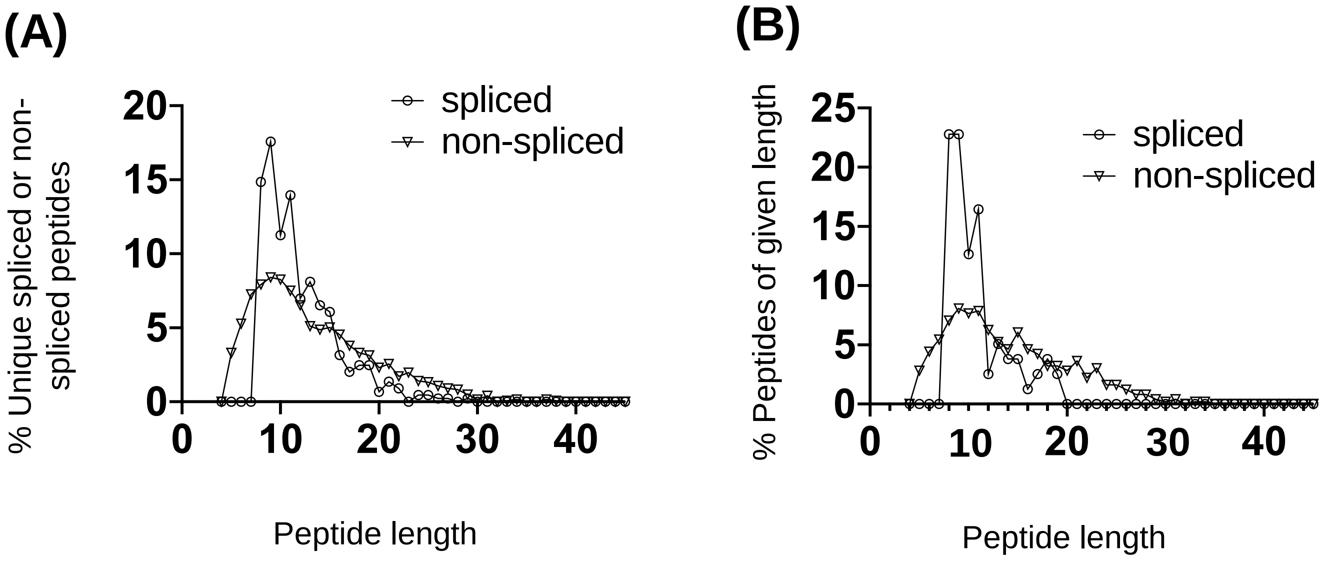


**Supplementary Figure 2. Length distribution of all peptide products generated from 2h *in vitro* proteasomal digests**

(A). Length distributions of all unique spliced (n=444) and non-spliced (n=1,212) peptide products generated from all 25 precursor polypeptide substrates after a 2h *in vitro* digest.

(B). Length distributions of unique spliced (n=156) and non-spliced (n=610) peptides generated from the 5 longest precursor polypeptide substrates (30-47 aa in length) after a 2 h *in vitro* digest.

### Supplementary Figure 3

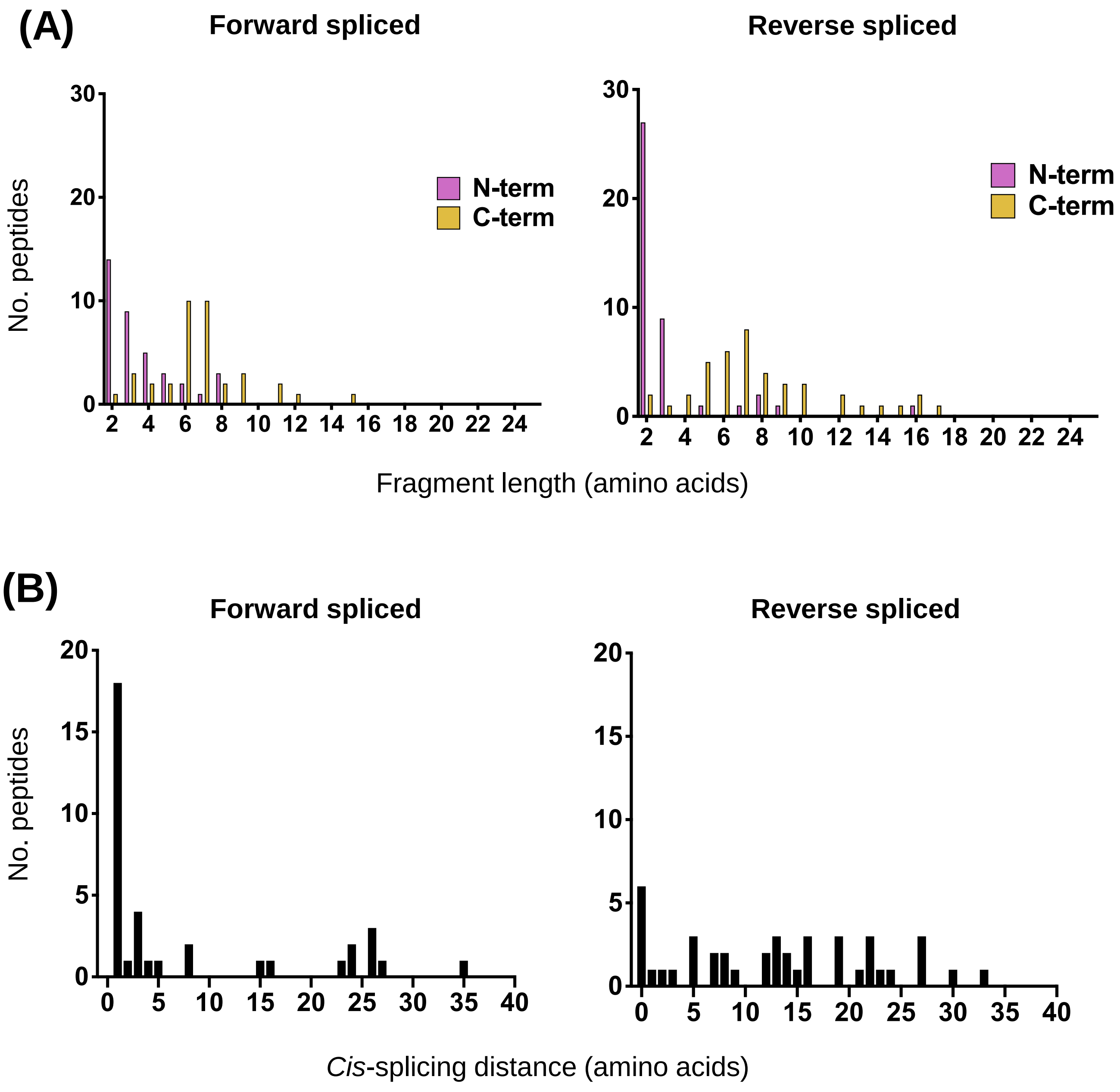


**Supplementary Figure 3. Fragment length and *cis*-splicing distributions for forward and reverse PCPS reactions occurring during digestion of the 5 longest polypeptide precursors**

(A) Fragment length distributions of N-terminal and C-terminal splice partners involved in forward and reverse PCPS reactions generating spliced peptides from the 5 longest polypeptide precursors (30-47aa in length).

(B). *Cis*-splicing distances in forward and reverse PCPS reactions generating spliced peptides from the 5 longest polypeptide precursors (30-47aa in length).
